# Clinical And Analytical Performance Of An Automated Serological Test That Identifies S1/S2 Neutralizing IgG In Covid-19 Patients Semiquantitatively

**DOI:** 10.1101/2020.05.19.105445

**Authors:** Fabrizio Bonelli, Antonella Sarasini, Claudia Zierold, Mariella Calleri, Alice Bonetti, Chiara Vismara, Frank Blocki, Luca Pallavicini, Alberto Chinali, Daniela Campisi, Elena Percivalle, Anna Pia DiNapoli, Carlo Federico Perno, Fausto Baldanti

**Affiliations:** DiaSorin, S.p.A., Saluggia, Italy; Molecular Virology Unit, Fondazione Istituto di Ricovero e Cura a Carattere Scientifico (IRCCS) Policlinico San Matteo, Pavia, Italy; Department of Clinical, Surgical Diagnostic and Pediatric Sciences, University of Pavia, Pavia, Italy; Department of Laboratory Medicine, ASST Niguarda Hospital, University of Milan, Milan, Italy

## Abstract

**BACKGROUND:** In the Covid-19 pandemic, highly selective serological testing is essential to define exposure to SARS-CoV-2 virus. Many tests have been developed, yet with variable speed to first result, and of unknown quality, particularly when considering the prediction of neutralizing capacity.

**OBJECTIVES/METHODS:** The LIAISON^®^ SARS-CoV-2 S1/S2 IgG assay was designed to measure antibodies against the SARS-CoV-2 native S1/S2 proteins in a standardized automated chemiluminescent assay. Clinical and analytical performance of the test were validated in an observational study using residual samples (>1500) with positive or negative Covid-19 diagnosis.

**RESULTS:** The LIAISON^®^ SARS-CoV-2 S1/S2 IgG assay proved highly selective and specific, and offers semiquantitative measures of serum or plasma levels of anti-S1/S2 IgG with neutralizing activity. The diagnostic sensitivity was 91.3% and 95.7% at >5 or ≥15 days from diagnosis respectively, and 100% when assessed against a neutralizing assay. The specificity ranged between 97% and 98.5%. The average imprecision of the assay was <5 % coefficient of variation. Assay performance at 2 different cut-offs was evaluated to optimize predictive values in settings with different % disease prevalence. CONCLUSIONS. The automated LIAISON^®^ SARS-CoV-2 S1/S2 IgG assay brings efficient, sensitive, specific, and precise serological testing to the laboratory, with the capacity to test large amounts of samples per day: first results are available within 35 minutes with a throughput of 170 tests/hour. The test also provides a semiquantitative measure to identify samples with neutralizing antibodies, useful also for a large scale screening of convalescent plasma for safe therapeutic use.

**IMPORTANCE:** With the worldwide advance of the COVID-19 pandemic, efficient, reliable and accessible diagnostic tools are needed to support public health officials and healthcare providers in their efforts to deliver optimal medical care, and articulate sound demographic policy. DiaSorin has developed an automated serology based assay for the measurement of IgG specific to SARS CoV-2 Spike protein, and tested its clinical performance in collaboration with Italian health care professionals who provided access to large numbers of samples from infected and non-infected individuals. The assay delivers excellent sensitivity and specificity, and is able to identify samples with high levels of neutralizing antibodies. This will provide guidance in assessing the true immune status of subjects, as well as meeting the pressing need to screen donors for high titer convalescent sera for subsequent therapeutic and prophylactic use.

## INTRODUCTION

SARS-CoV-2, the virus responsible for the Covid-19 pandemic, has spread at an alarming rate since the first case tracked back to mid-November of 2019 in Wuhan China (1). Contraction and subsequent transmission accrues most prevalently from community exposure, non-human exposure, or amongst relatives living in proximity to symptomatic or asymptomatic infected individuals. Due to the lack of readily available diagnostics, inferior means of infection control, or the inability to triage and isolate both acute and suspected cases consequent to space limitations, the Covid-19 pandemic, in placing increasingly excessive demands on the global healthcare network, has unveiled a number of critical limitations (2).

No safe vaccines have been developed for SARS-CoV infections to date, and the lack of currently available effective antiviral therapies, in spite of years of ongoing research, are certainly hampering efforts to combat this pandemic. In addition, the current molecular-based diagnostic tools utilized to diagnose infection, though serving adequately as the only means available, are not suitable for mass screening, and though many serological assays have been developed, no scientific data are available to authenticate their effectiveness. Finally, once the World Health Organization (WHO) recognized the SARS-CoV-2 outbreak as a Public Health Emergency of International Concern on January 30, 2020, efforts have been hampered internationally, nationally, and locally by a lack of coordinated guidance to properly inform public policy makers, and a lack of ready access to accurate and rapid testing.

Understanding the efficiency of community transmission of SARS-CoV-2, including the contribution of mild or asymptomatic cases, represents a knowledge gap (1). As of May 5, 2020, global cases have surpassed 3 million, with 254,592 registered deaths (3). Global mortality consequent to SARS-CoV-2 infection is running at 7.0% with national rates for France, Italy, Spain, United Kingdom, Germany, and the United States running at 14.9%, 13.7%, 11.7%, 15.0%, 4.2%, and 5.8% (4).

In view of these daunting numbers, effective, sensitive and specific means for the identification and laboratory confirmation SARS-CoV-2 infection are urgently needed. In response to these needs, DiaSorin has developed a highly sensitive, specific, automated and contained chemiluminescence serological assay for the detection of SARS-CoV-2 Spike protein-specific antibodies with neutralizing potential from serum or plasma, to be used in diagnostic, epidemiological and vaccine evaluation studies.

Specifically, it is envisioned that the test may be used to: 1) screen infected health care workers and the general population for recovery and/or past exposure; 2) epidemiological studies characterizing the demographics of viral spread and the efficacy of containment measures directed towards SARS-CoV-2 at the local, national, and international level; 3) screen convalescent sera for both therapeutic and prophylactic use, and 4) evaluate vaccine effectiveness in clinical studies.

## RESULTS

### Sample characteristics

Clinical assessment of the LIAISON^®^ SARS-CoV-2 S1/S2 IgG assay was performed using various sample groups (Figure 1). All the patient groups categorized as positive for Covid-19 were significantly different from the negative groups at P<0.0001 as determined by a pairwise t-test comparison with Bonferroni multiplicity adjustment. Median S1/S2 IgG levels were 96.3 AU/mL (95% CI 85.8 to 108.0 AU/mL, N=64), 28.6 AU/mL (95% CI 10.6 to 45.1 AU/mL, N=67), and 15.5 AU/mL (95% CI 5.7 to 32.2 AU/mL, N=80) for intensive care unit (ICU) patients, hospitalized patients, and RT-PCR-positive patients, respectively. Median levels of negative samples were 2.3 AU/mL (95% CI 2.2 to 2.4 AU/mL, N=1140), 2.4 AU/mL (95% CI 2.1 to 2.9 AU/mL, N=50), and 2.2 AU/mL (95% CI 1.8 to 4.6 AU/mL, N=10) for pre-Covid-19, RT-PCR negative, and other coronavirus (non-SARS-Cov-2) subjects, respectively. In addition, the ICU patient group had significantly higher levels of S1/S2 IgG compared to hospitalized patient group (P<0.0001). Table 1 further dissects the temporal component of distribution with early samples having low levels of S1/S2 IgG presenting a low positive predictive agreement to RT-PCR at time ≤5 days from diagnosis (33.3%), increasing to 95.7% at ≥15 days from diagnosis.

**Table 1:** Longitudinal Assessment of Positive Predictive Agreement (PPA) to RT-PCR Diagnosis in Covid-19 Patients. Serial samples from 104 Covid-19 patients positive by RT-PCR admitted to the hospital or ICU were tested with the LIAISON^®^ SARS-CoV-2 S1/S2 IgG assay. A value of 9 AU/mL was used as the cut-off for positivity.

| Days from<br>Diagnosis | First Serial<br>Measurement |  | Second Serial<br>Measurement |  | Third Serial<br>Measurement |  | Cumulative | PPA (95%CI) |
| --- | --- | --- | --- | --- | --- | --- | --- | --- |
|  | Total<br>N | S1/S2<br>IgG<sup>+</sup> | Total<br>N | S1/S2<br>IgG<sup>+</sup> | Total<br>N | S1/S2<br>IgG<sup>+</sup> | S1/S2<br>IgG<sup>+</sup>/Total |  |
| ≤ 5 days | 84 | 28 |  |  |  |  | 28/84 | 33.3%<br>(23.4% to 44.5%) |
| 6-14 days | 7 | 7 | 71 | 62 | 1 | 1 | 70/79 | 88.6%<br>(79.5% to 94.7%) |
| ≥15 days | 13 | 13 | 12 | 12 | 22 | 20 | 45/47 | 95.7%<br>(85.5% to 99.5%) |
| **Total**<br>**Subjects** | 104 |  | 83 |  | 23 |  |  |  |

**Figure 1:**
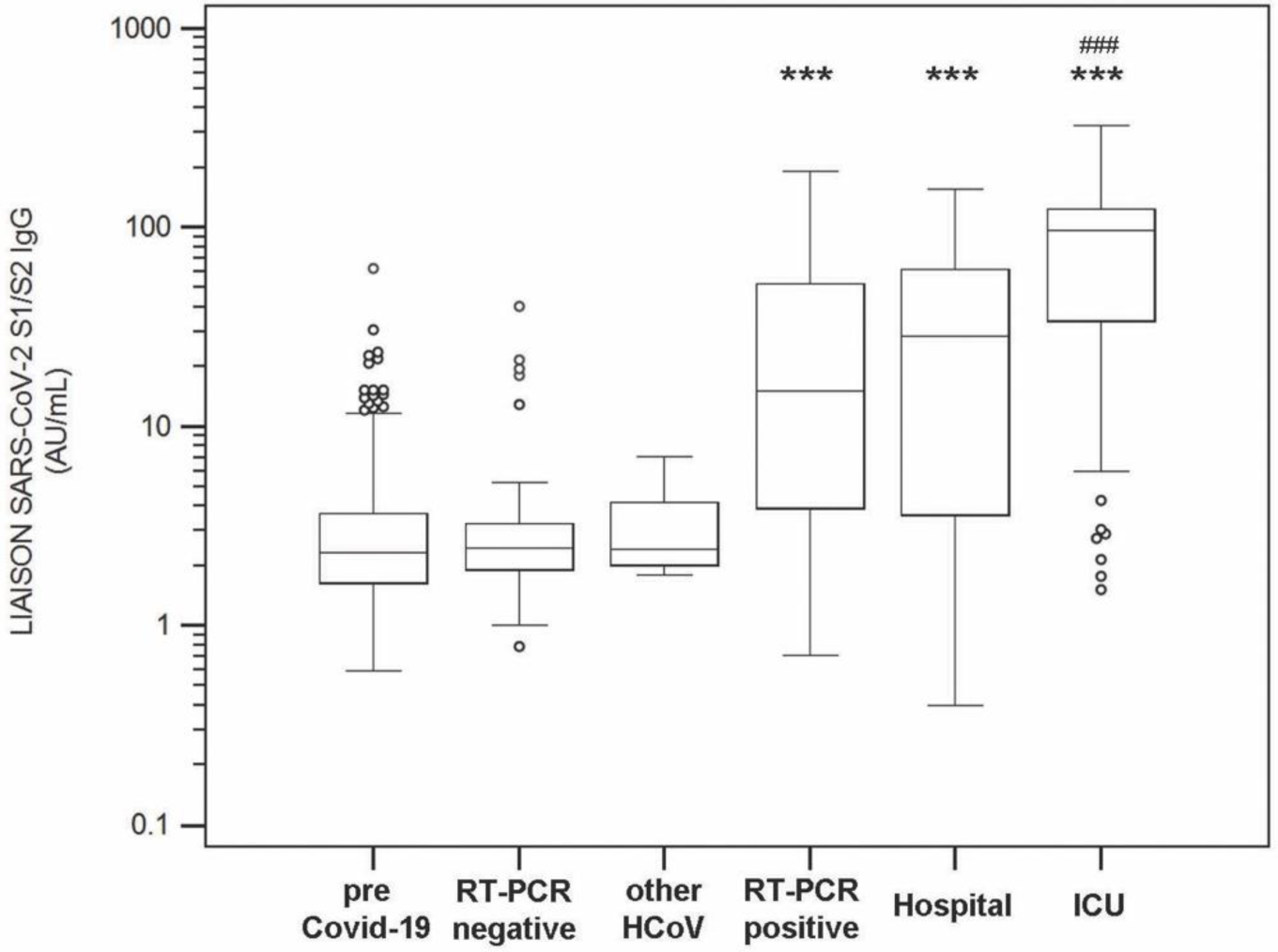
Distribution of SARS-CoV-2 S1/S2 IgG levels in various patient groups. All the patient groups categorized as positive (RT-PCR-positive, hospitalized and ICU) are significantly different from the negative groups (RT-PCR-negative, pre-Covid-19 and other HCov) at P<0.0001 (***), as determined by a pairwise t-test comparison with Bonferroni multiplicity adjustment. The ICU patient group is significantly different from the hospitalized patient group (p<0.0001, ###)

### Clinical Performance

A receiver operating characteristic analysis was fitted to determine the best cut point supporting positive diagnoses. The maximum Youden index occurs at a cut point of 9.4 (sensitivity / specificity of 95% / 97% respectively), with an area under the curve of 0.980 (95% CI 0.960-0.990, p<0.0001, Figure 2).

**Figure 2:**
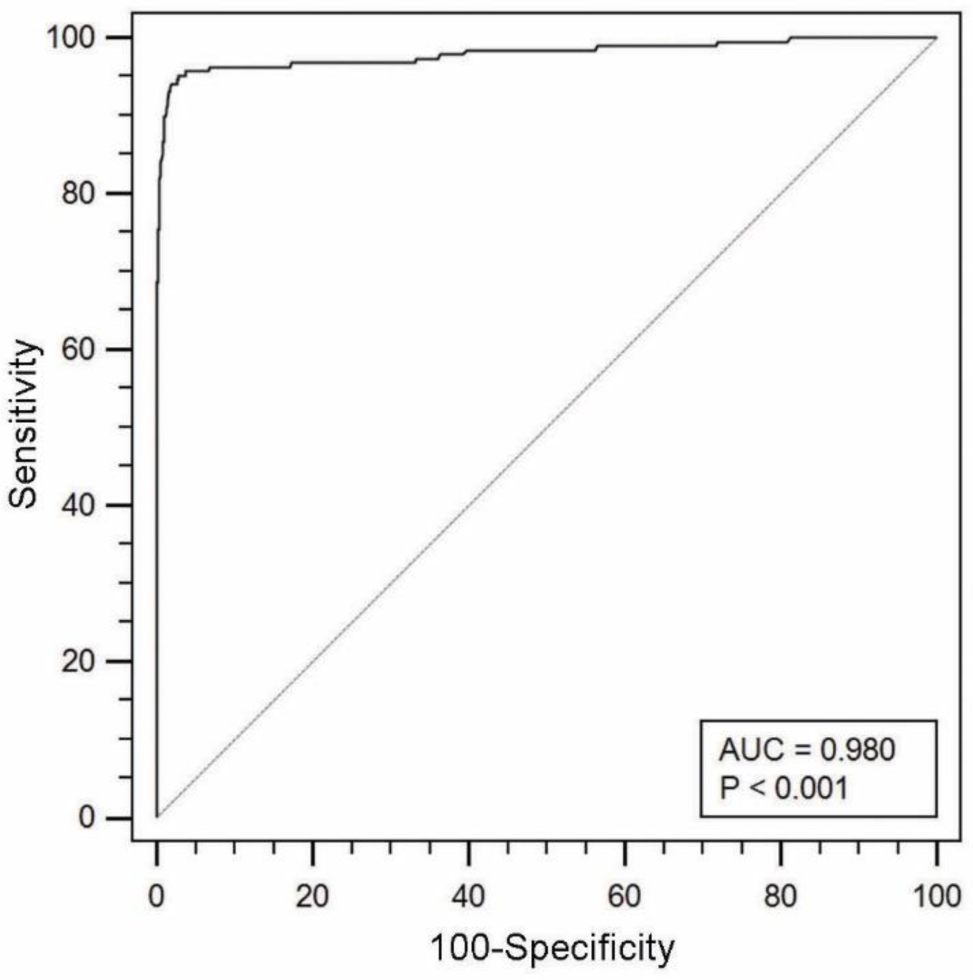
Receiver operating curve for distinguishing samples from patients affected by Covid-19 using the SARS-CoV-2 S1/S2 IgG test in a group of 1568 samples (N=188 positives). Area under the curve, AUC=0.980 (95% CI 0.960-0.990). The Youden Index associated cut-off is 9.4 AU/mL (95% CI >7.1 AU/mL to >12.1 AU/mL).

Clinical performance of the LIAISON^®^ SARS-CoV-2 S1/S2 IgG assay is shown in Table 2. The sensitivity was determined by investigating 211 samples collected longitudinally over the course of time from 84 patients at admission and thereafter variably up to 36 days. Infection with SARS-CoV-2 was confirmed by a positive RT-PCR test at the early phase of infection at time of diagnosis. Logarithmic values of SARS-Cov-2 S1/S2 IgG are plotted over time with a fitted curve (Figure 3), projecting estimations of 5 days for an average sample to reach 15 AU/mL, with 92.9% of the samples exceeding the 15 AU/mL threshold by 5 or more days post diagnosis. Table 2 compares the sensitivities and specificities consequent to higher cut-off of 15 AU/mL as currently suggested by the manufacturer’s instructions for use. Diagnostic sensitivity with the lower cut-off is calculated at 33.3% for the early samples (≤ 5 days after diagnosis) and 91.3% for samples collected >5 days post diagnosis. Diagnostic sensitivity with the higher cut-off drops to 22.6%% for the early samples (≤ 5 days after diagnosis) and 88.2% for samples collected >5 days post diagnosis. Conversely, specificity (from the testing of 1140 stored residual samples from laboratory routine collected before the Covid-19 outbreak) increased slightly from 97.1% to 98.5% at the lower and higher cut-offs, respectively. Specificities evaluated using all negative samples were 97.0% and 98.1%. The importance of choosing a cut-off that provides higher sensitivity (9 AU/mL) versus one that provides lower sensitivity but higher specificity (15 Au/mL) is influenced by the disease prevalence, reflected in positive and negative predictive values (PPV and NPV). When the intent is to use the assay for screening, a higher threshold may be desirable, whereas, in a high prevalence environment such as hospitals caring for high numbers of Covid-19 subjects, when the test is used for aid in diagnosis, the lower threshold 9 AU/mL may be preferred (Table 3).

**Table 2:**
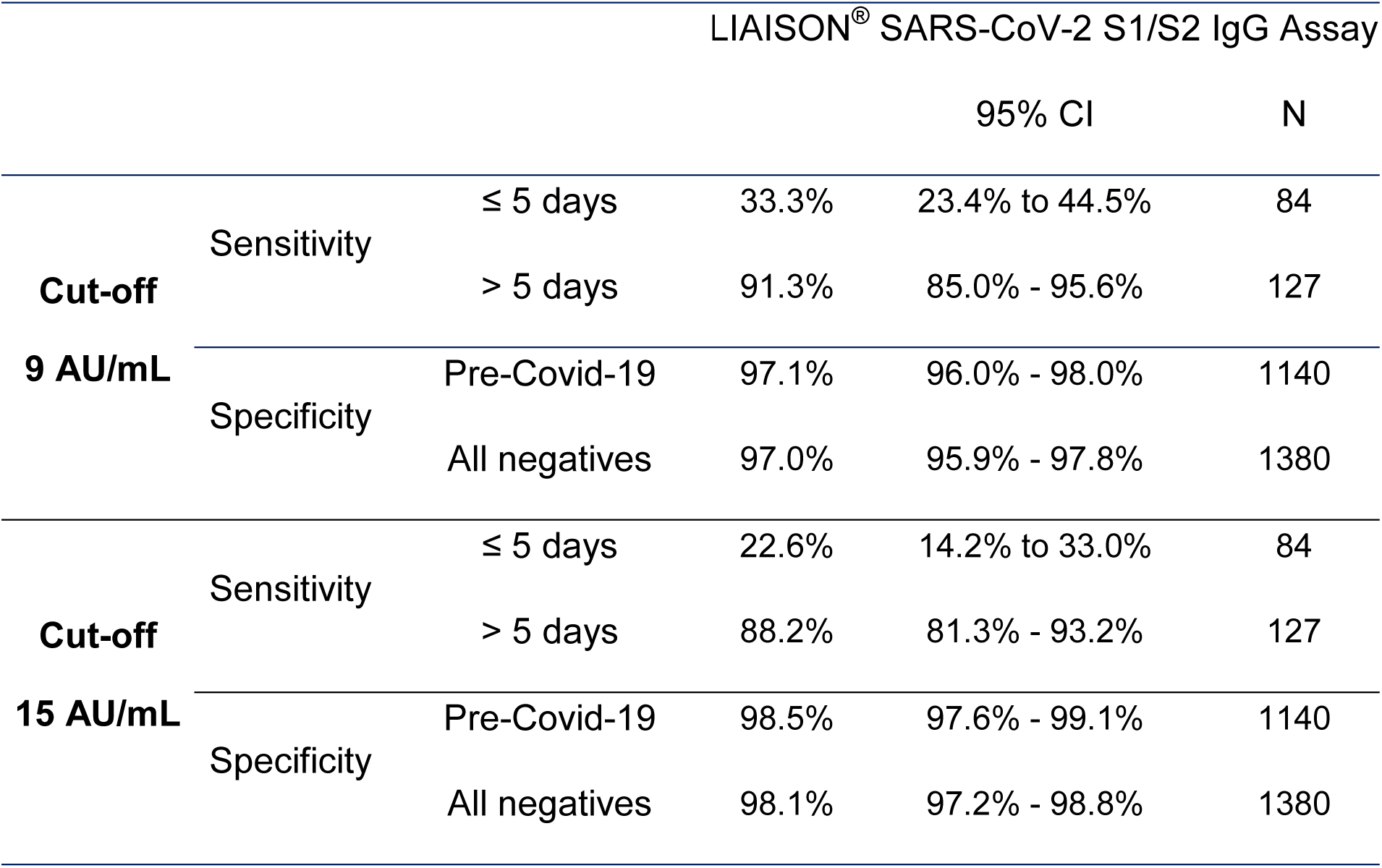
Clinical Performance of the LIAISON^®^ SARS-CoV-2 S1/S2 IgG Assay Using 9 and 15 AU/mL as Cut-off Based on RT-PCR Diagnoses.

**Table 3:**
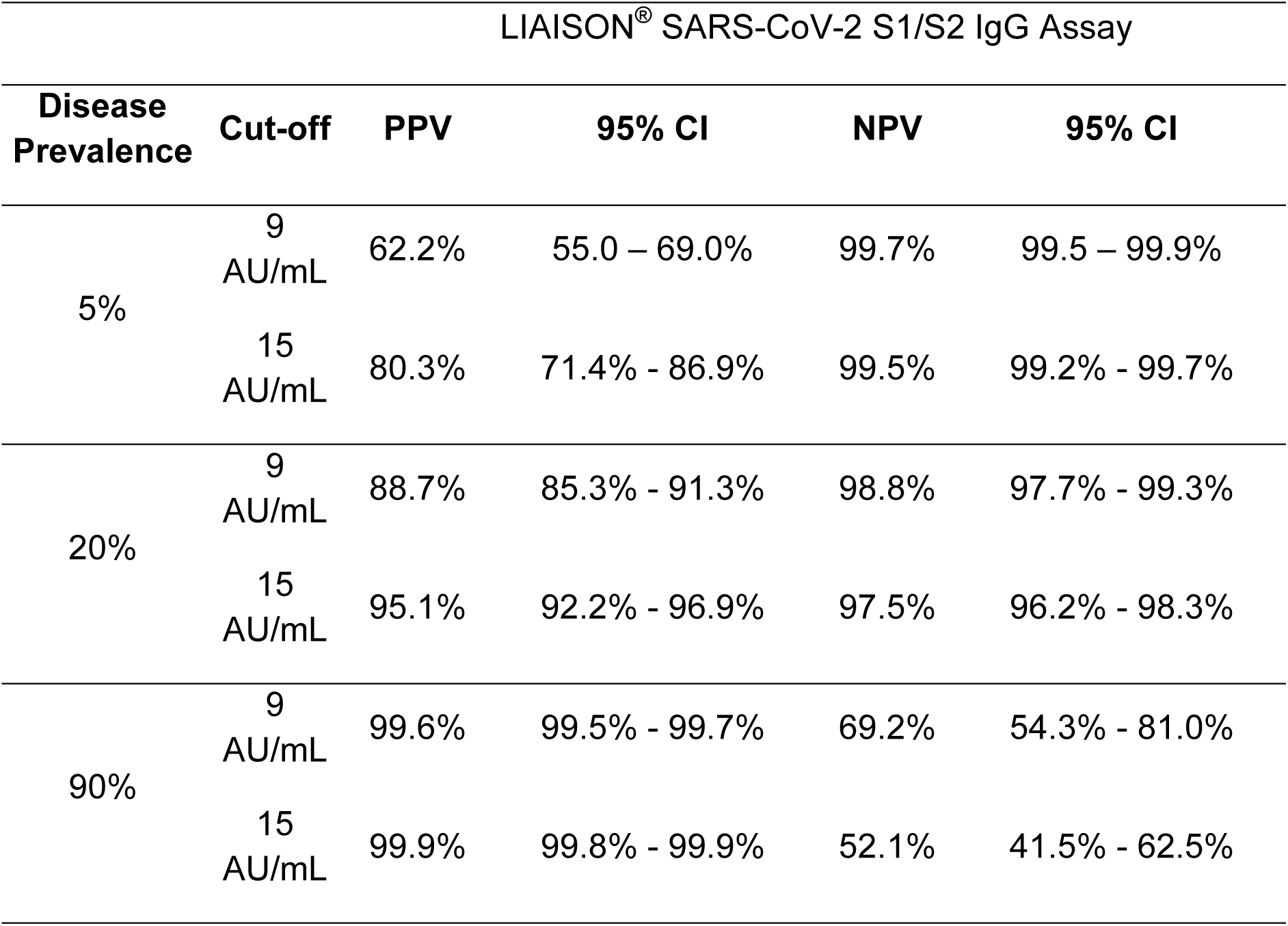
Positive And Negative Predictive Values at Various Disease Prevalence Thresholds for the LIAISON^®^ SARS-CoV-2 S1/S2 IgG Assay at Cut-offs of 9 and 15 AU/mL

**Figure 3:**
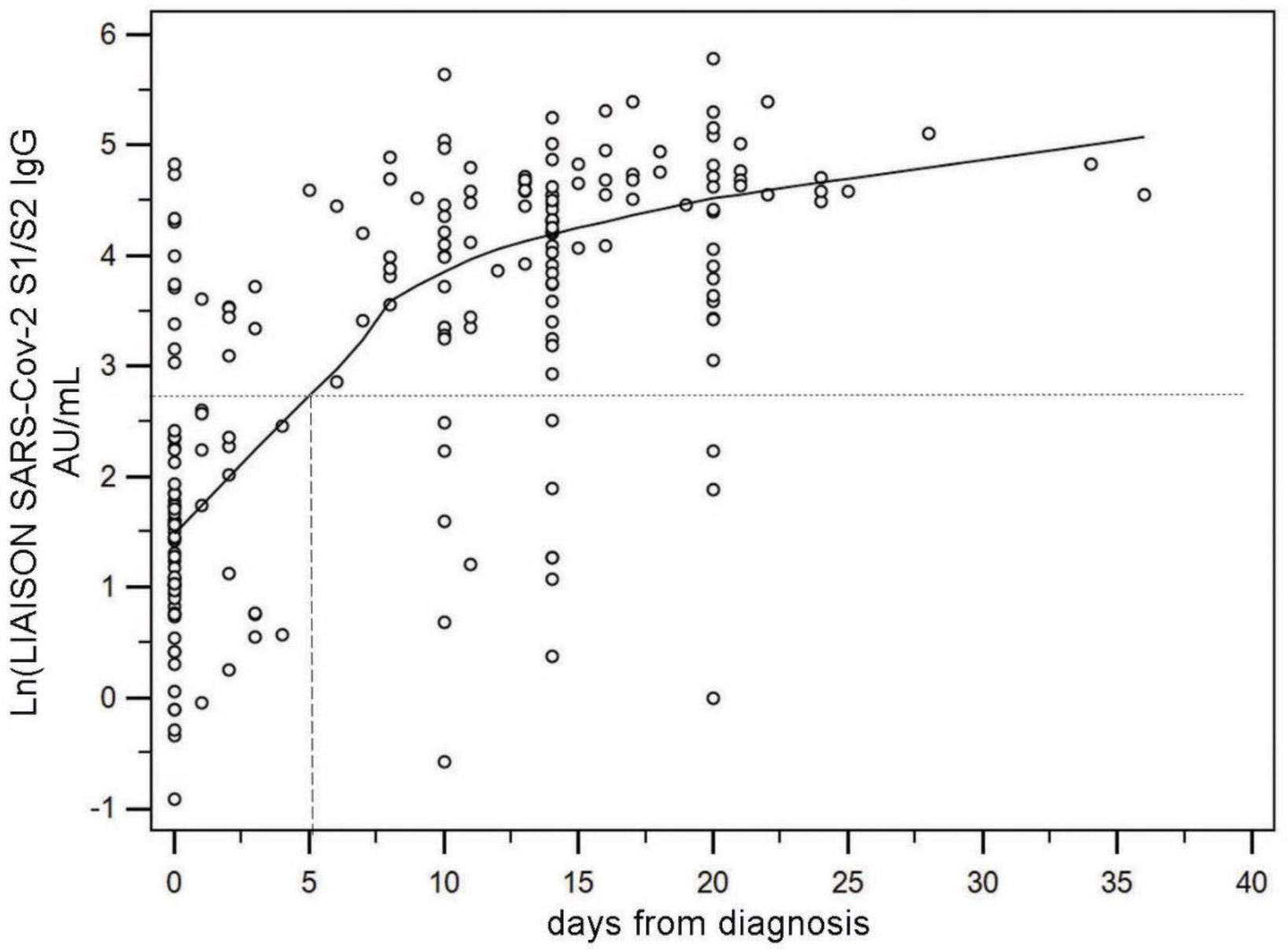
SARS-CoV-2 S1/S2 IgG measurements for 211 samples collected over the course of time from 84 patients at admission and variably thereafter up to 36 days. The upward trend was modeled using an exponential regression ln(IgG)=A+B* exp(C*Days). The parameter A (4.58) represents the upper limit to which the LIAISON^®^ SARS-CoV-2 S1/S2 IgG trends over time and corresponds to 98 AU/mL. A+B (1.63) is the value at time zero corresponding to 5.1 AU/mL on the original scale. The parameter C (−xs0.112) is the rate at which the curve moves up to the asymptote and corresponds to 1.1 AU/mL per day. The dotted horizontal line is 15 AU/mL, the positive cut-off of the assay. The curve cuts this at 5 days, giving an estimate of the delay from diagnosis above which the patient is on average positive.

### Comparison to samples with neutralizing titer

Comparison to a neutralization (NT) assay was evaluated by testing 304 samples collected during the outbreak from subjects whose NT assay results were available: 180 were NT assay-negative, and 124 were NT assay-positive (titer >1:40). Positive agreement was 94.4% (95% CI 88.8% - 97.2%) and negative agreement was 97.8% (95% CI 94.1% - 99.1%). The relationship between the LIAISON^®^ SARS-CoV-2 S1/S2 IgG assay and NT assay-negative or NT assay-positive samples portrays a nearly complete separation between the 2 groups with medians of 2.4 AU/mL (95% CI 2.2 to 2.6 AU/mL), and 61.8 AU/mL (95% CI 50.3 to 70.7 AU/mL), respectively (Figure 4). In Figure 5A, the LIAISON^®^ SARS-CoV-2 S1/S2 IgG assay’s measurements were separated into 3 semi-quantitative groups (<40 AU/mL, 40-80 AU/mL, and >80 AU/mL) and related to NT assay titers ≥1:160, which is the threshold recommended by the FDA guidelines for use in convalescent blood transfusion. 39% (17/43), 56% (24/43), and 87% (33/38) of the samples, respectively, had NT assay titers ≥1:160 (5). Furthermore, since the FDA guidelines also admit NT assay titers of ≥1:80 as acceptable, additional leeway is granted towards use of the LIAISON^®^ SARS-CoV-2 S1/S2 IgG assay to pre-screen or assess blood donor samples for potential convalescent plasma/serum therapy: 92% (35/38) and 79% (34/43) of the > 80 AU/mL and 40-80 AU/mL groups, respectively, had NT assay titers ≥1:80 (Figure 5B).

**Figure 4:**
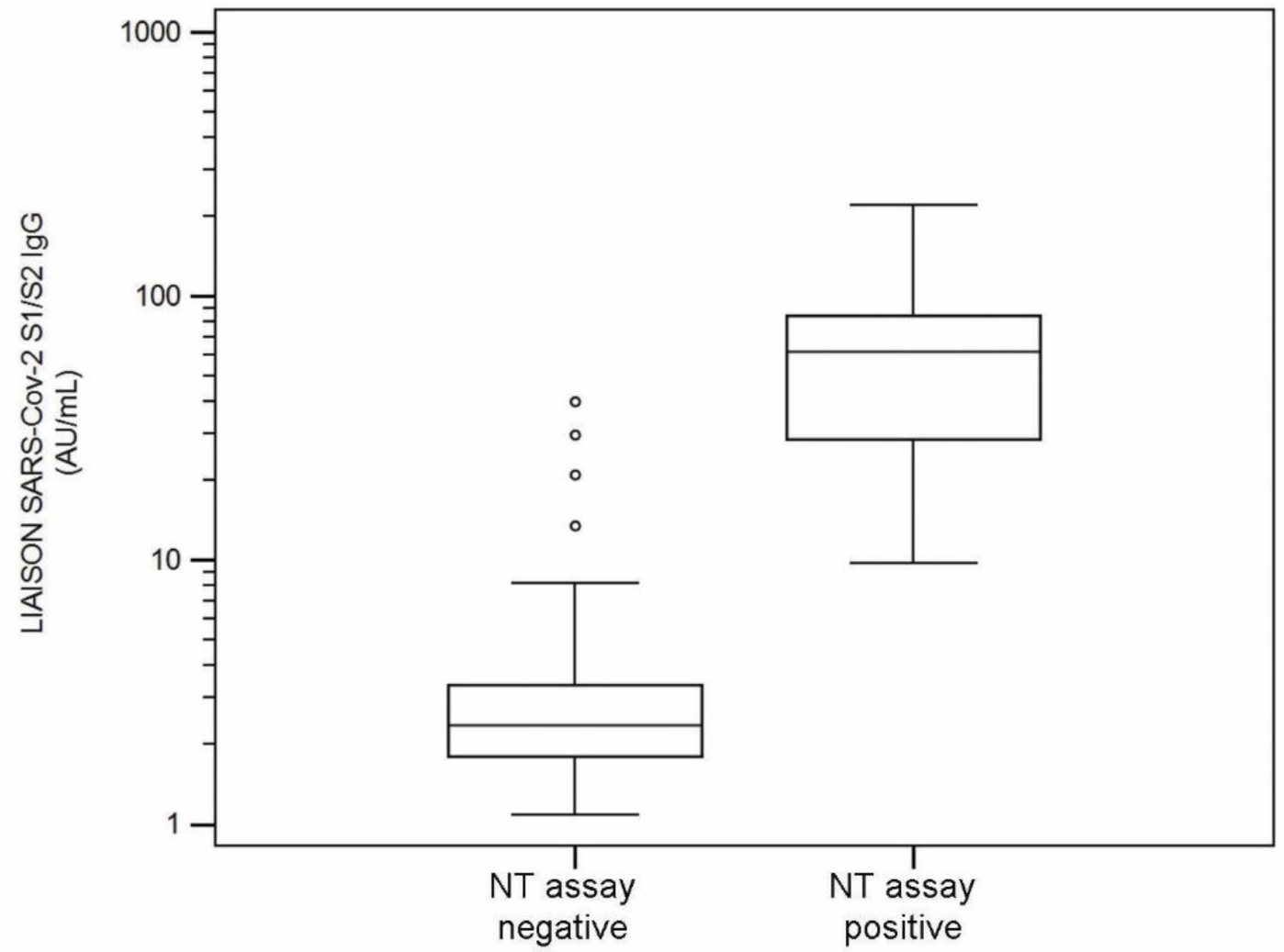
Distribution of the LIAISON^®^ SARS-CoV-2 S1/S2 IgG assay measurements compared to positive and negative samples by the NT assay.

**Figure 5:**
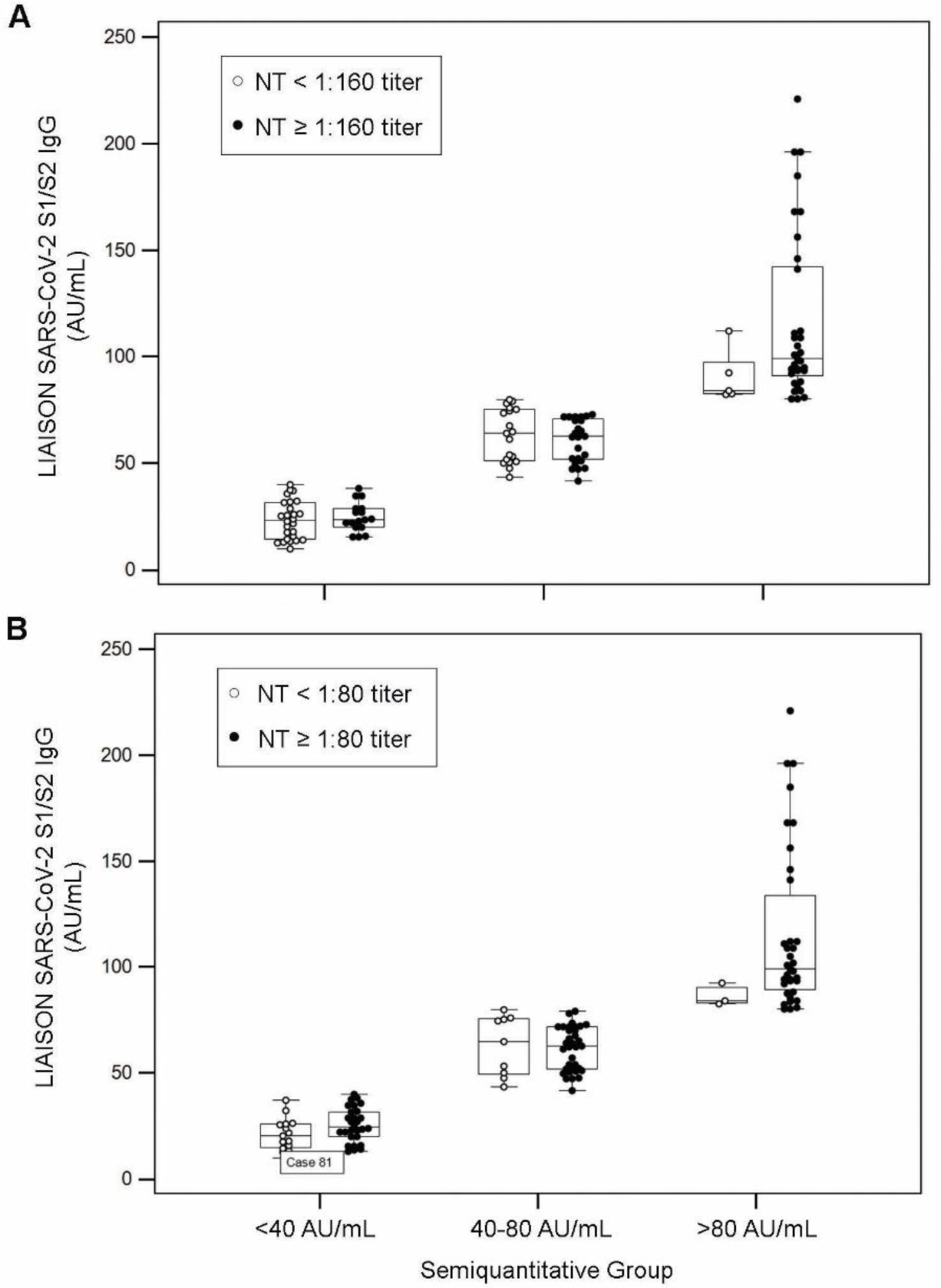
Relationship and distribution of the LIAISON^®^ SARS-CoV-2 S1/S2 IgG assay levels versus NT dilutions. LIAISON^®^ SARS-CoV-2 S1/S2 IgG measurements were separated into 3 groups (<40 AU/mL, 40-80 AU/mL, and >80 AU/mL) and related to NT assay grouped by titer ≥ 1:160 **(A)** or ≥ 1:80 **(B)**. 39% (17/43), 56% (24/43), and 87% (33/38) of samples have a NT assay titer ≥1:160, while 65% (28/43), 79% (34/43), and 92% (35/38) of samples have a NT assay titer ≥1:80. Both titers are considered acceptable by FDA guidelines (5).

### Analytical Performance

The LIAISON^®^ SARS-CoV-2 S1/S2 assay was evaluated for intra-assay imprecision using 6 samples with moderate, low, or negative S1/S2 IgG levels. The average intra-assay imprecision was 2.8 %CV (range 2.0-3.4%CV), and total-assay imprecision averaged 3.2%CV (range 2.7-3.9%CV) (Table 5).

**Table 4:**
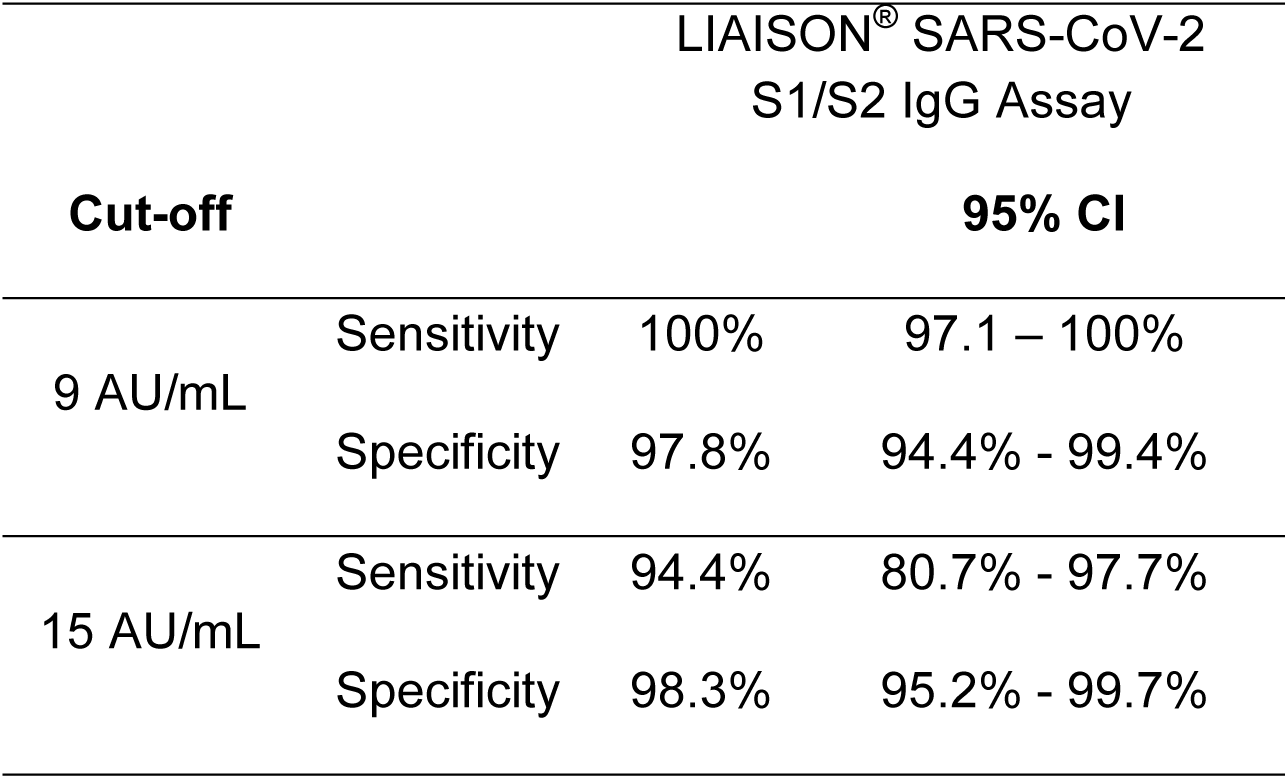
Comparison of the LIAISON^®^ SARS-CoV-2 S1/S2 IgG Assay Measurement Using 9 and 15 AU/mL as Cut-offs from Samples with Known Neutralizing Titers Defined as Negative (< 1:40) or Positive for Neutralizing Antibodies (≥ 1:40).

**Table 5:** Imprecision Data of the LIAISON^®^ SARS-CoV-2 S1/S2 Assay from a 5 Day Precision Study Conducted According to CLSI EP5-A3 Guidelines. The Panel Samples Were Tested in 6 Replicates per Run, 3 Runs per Day for 5 Operating Days.

| Sample | N | Mean<br>(AU/mL) | Intra |  | Total |  |
| --- | --- | --- | --- | --- | --- | --- |
|  |  |  | SD | CV % | SD | CV % |
| Negative 1 | 90 | 5.45 | 0.14 | 2.5 | 0.15 | 2.7 |
| Negative 2 | 90 | 6.72 | 0.23 | 3.4 | 0.26 | 3.9 |
| Low 1 | 90 | 11.1 | 0.34 | 3.0 | 0.38 | 3.4 |
| Low 2 | 90 | 20.2 | 0.54 | 2.7 | 0.67 | 3.3 |
| Moderate 1 | 90 | 40.1 | 1.14 | 2.8 | 1.17 | 2.9 |
| Moderate 2 | 90 | 64.1 | 1.68 | 2.6 | 1.86 | 2.9 |

Cross-reactivity with other coronaviruses was tested against 10 patient samples positive by their respective RT-PCR tests to other coronaviruses that maintained negative NT assay results by SARS-Cov-2. Their LIAISON values ranged from 1.81 to 7.09 AU/mL, with an average of 3.45 AU/mL which falls far below both cut points of 9 and 15 AU/mL indicating the absence of cross-reactivity with the other coronaviruses tested (Table 6). Additionally, cross-reactivity was assessed in samples from patients with conditions caused by other viruses, other organisms, or with atypical immune system activity. As shown in Table 7, 3 out of 160 assessed specimens (1.9%) resulted positive with the LIAISON^®^ SARS-CoV-2 S1/S2 IgG assay. Potentially interfering substances such as triglycerides (3000 mg/dL), cholesterol (400 mg/dL), hemoglobin (1000 mg/dL) conjugated and unconjugated bilirubin (40 mg/dL), acetaminophen (500 mg/mL and ibuprofen (500 mg/mL) showed no interference at the indicated concentrations. The LIAISON^®^ SARS-CoV-2 S1/S2 IgG assay demonstrated a negative bias up to a 16% in S1/S2 IgG-positive specimens with biotin concentrations above 3500 ng/mL, a concentration 15-fold higher than that induced following ingestion of a 20 mg/day biotin supplement (6).

**Table 6:**
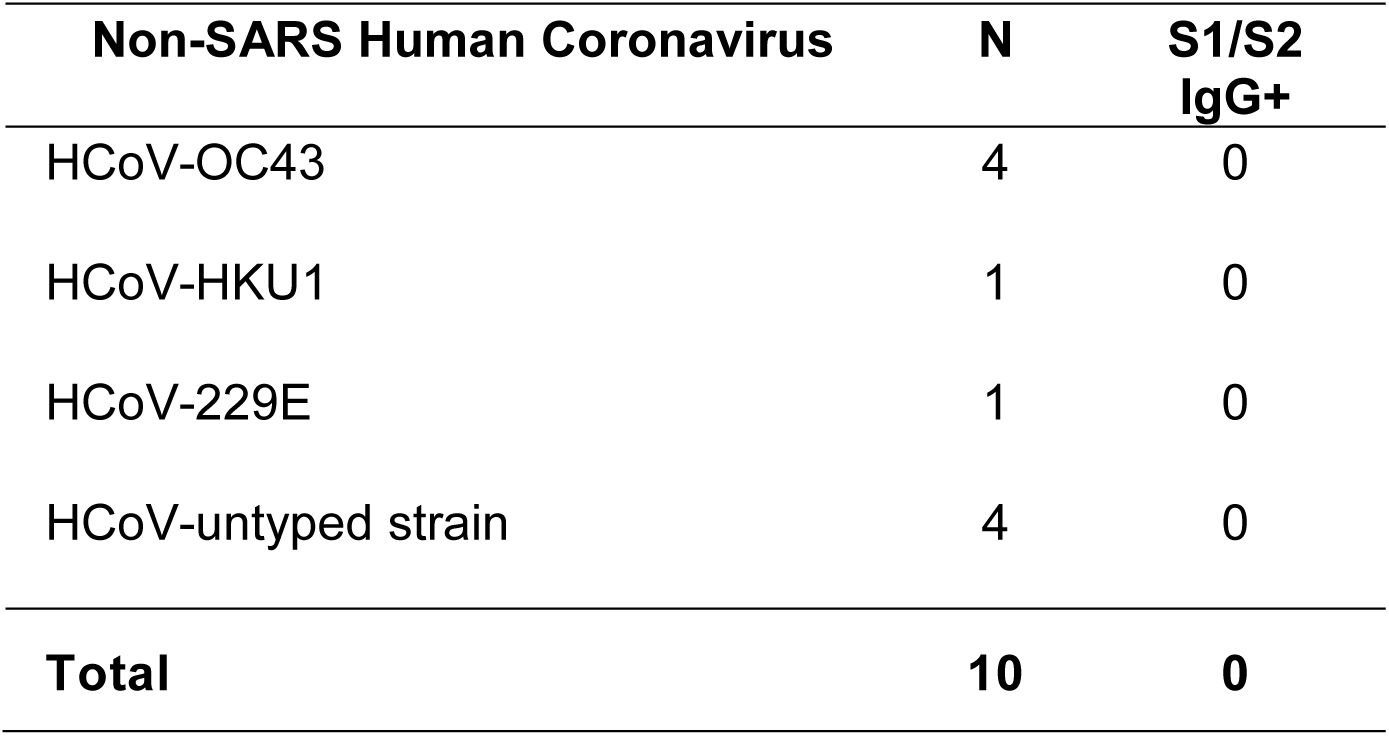
Cross-reactivity with Other Coronaviruses Tested in Patient Samples Positive by their Respective RT-PCR Tests.

**Table 7:** Cross-reactivity with Other Conditions Caused by Other Viruses, Other Organisms, or with Atypical Immune System Activity with Symptoms Similar to Covid-19.

| Condition | N | S1/S2<br>IgG<sup>+</sup> |
| --- | --- | --- |
| Nuclear autoantibodies (ANA) | 10 | 0 |
| HBV | 10 | 1 |
| HCV | 10 | 0 |
| Influenza A | 10 | 1 |
| Influenza B | 10 | 0 |
| Respiratory syncytial virus | 10 | 0 |
| *Borrelia burgdorferi* | 10 | 0 |
| *Mycoplasma pneumoniae* | 10 | 0 |
| EBV | 10 | 0 |
| CMV | 10 | 0 |
| HSV-1/2 | 10 | 0 |
| HAMA | 10 | 0 |
| Parvovirus B19 | 10 | 0 |
| Rheumatoid factor | 10 | 1 |
| Rubella | 10 | 0 |
| VZV | 10 | 0 |
| **Total** | **160** | **3** |

**ABBREVIATIONS**
ICU: intensive care unit

## DISCUSSION

SARS-Cov-2 is a single strand, positive sense RNA virus that is most closely related to SARS-CoV and other B lineage members of the β genogroup of coronaviruses (7, 8). Other readily recognized members of the coronavirus family include the extremely virulent MERS CoV, and the less virulent OC43, HKU1, 229E and NL63, HCoVs more commonly associated with the common cold in adults. Coronavirus RNA encodes for four major categories of structural protein including Membrane, Envelope, Nucleocapsid and Spike that are referred to as M, E, N and S, respectively. While N protein elicits cell mediated immunity attributable to two predominant CD8 T cell epitopes (9), of the remaining three structural proteins, S protein is widely recognized as that most specific with regard to generating protective, neutralizing antibodies (10, 11).

The specificities reported from *in vitro* diagnostic immunoassays (12-14) are impacted greatly by the fidelity of preservation of both linear and conformational epitopes of the given analyte being measured for presentation to specific immunoglobulins within a patient’s serum sample (here SARS-Cov-2 S1/S2). Specificity is frequently significantly compromised by the casual manner in which most ELISA assays are fabricated. This is consequent to the generally accepted means of passive adsorption of the target analyte protein to the plastic or nylon surface of microtiter plates. This procedure induces significant structural deformation and denaturation, with the consequent loss of native conformation, as well as occlusion of access to both conformational and linear epitopes beneath the protein stuck to the plastic titer plate’s surface. As explained in Materials and Methods, our system allows for optimal maintenance of Spike protein conformation. Consequently, the LIAISON^®^ SARS-CoV-2 S1/S2 IgG assay rendered no false positive results from NT assay-negative, RT-PCR-positive samples for related coronavirus members, and its performance is sensitive, specific, and precise as evaluated in >1500 samples.

The use of convalescent serum to treat subjects with acute SARS-Cov-2 has growing appeal for meeting the immediate challenges being imposed upon increasingly stressed health care systems, in light of the premature status of vaccines, and limited availability of effective anti-viral therapeutic regimens (15-19). The LIAISON^®^ SARS-CoV-2 S1/S2 IgG assay was designed to detect IgG with neutralizing potential, and is shown here to have very good sensitivity and specificity in identifying samples with positive neutralization titers. Furthermore, if used in a semi-quantitative manner, higher LIAISON units are indicative of higher NT assay titers, and provide a pre-screen tool to assess large numbers of samples. While neutralization tests provide the recognized benchmark, they are not practical for implementation on a large scale screening basis, due to requirements for high biosecurity containment laboratories, and the need for highly trained personnel to execute labor-intensive protocols. With our system, clear separation of NT assay-negative samples from NT assay-positive samples was achieved. In fact, with 40-80 AU/mL levels measured by the LIAISON^®^ SARS-CoV-2 S1/S2 IgG assay, the probability to have neutralization titers ≥1:80 and ≥1:160 was 79% and 56%, while with >80 AU/ml the probability of having neutralization titers ≥1:80 and ≥1:160 was 92%, and 87%, respectively. This may be useful for the efficient screening of convalescent plasma for safe therapeutic use.

The LIAISON^®^ SARS-CoV-2 S1/S2 IgG assay’s sensitivity increases significantly as the immune response matures, as one would expect for an IgG-based serology assay’s assessment of a host response to viral infection (Table 1 & 2, and Figure 3). Here, sensitivities of 33.3% at <5 days but >91% at ≥5 days post admission on samples from 104 Italian patients whose RT-PCR tests were positive at the time of diagnosis are reported. Serology tests are now being utilized to gain an initial assessment of infection prevalence with reported numbers of ∼20% and ∼3% from New York and California, respectively (19). In California, a negative test with the LIAISON assay would have an accompanying NPV of >99.5%, and in NYC a NPV of >97.5%, regardless of cut-off, indicating that staying at home and avoiding exposures would be the best option. However, a positive test presents a quite different story, whereby California’s PPV of 80% derived from the higher cut-off, though overall less sensitive, would provide a positive test result, affording more confidence of the subject’s true positivity, while the lower cut-off would present some ambiguity as regards any subject’s real level of protection (PPV of 62%). In New York, however, regardless of the cut-off, the PPV of 89-95% affords a much greater degree of confidence that an individual would have protective levels of antibody. When testing in a hospital setting, the lower cut-off may be preferable to ensure a higher NPV, even though the overall specificity may be decreased.

In conclusion the automated LIAISON^®^ SARS-CoV-2 S1/S2 IgG assay brings efficient, sensitive, specific, and precise serological testing to the laboratory. Further, the assay is amenable for semi-quantitative efficient pre-screening of samples for neutralizing antibody content, to be used in convalescent plasma therapy.

## MATERIALS AND METHODS

### Assay format

The LIAISON^®^ SARS-CoV-2 S1/S2 IgG chemiluminescent assay is a recently developed assay from DiaSorin designed to detect IgG antibodies in the serum or plasma of subjects and patients exposed to the SARS-CoV-2 virus. The assay consists of paramagnetic microparticles (PMPs) coated with S1 and S2 fragments of the viral surface Spike protein (8). Recombinant fusion antigens were expressed in human cells (HEK-293) to ensure proper folding, oligomer formation and glycosylation, providing capture moieties more similar to the viral Spike proteins, as processed by natural cellular cleavage (20-22): this distinguishes the DiaSorin CLIA from commonly used ELISAs where the antigens are presented on plastic plate surfaces, and are susceptible to significant denaturation consequent to passive adsorption to these hydrophobic surfaces (23, 24). Distally biotinylated-S1 and biotinylated-S2 proteins were tethered to the surface of paramagnetic particles coated with streptavidin to assure optimal presentation of both S1 and S2 for access and recognition by specific immunoglobulin within pathologic serum samples.

The automated assay format consists of a first incubation step (10 minutes) of S1/S2-coated PMPs with patient sample (20 μl of either plasma or serum) in assay buffer to allow the binding of IgG in the sample specific to the antigens, followed by a wash step to remove unbound materials. Next ABEI (N-(4-aminobutyl)-N-ethyl-isoluminol)-labeled polyclonal goat anti-human IgG are added to the PMPs and further incubated for 8 minutes. After a final wash cycle, starter reagents are added and emitted relative light units (RLU) proportional to the sample’s anti-S1/S2 IgG levels are converted to arbitrary units (AU/mL) based on a standardized master curve. The automated assay is standardized based on a pool of patient samples with high S1/S2 IgG titers. First results are available within 35 minutes, and the throughput is 170 tests/hour.

### Analytical assay performance

A 5 day precision study according to CLSI EP5-A3 guidelines was performed using a panel of 6 plasma samples, prepared by either spiking or diluting as necessary to obtain negative, low positive and moderate positive samples. The panel samples were tested with LIAISON^®^ SARS-CoV-2 S1/S2 IgG assay in 6 replicates per run, 3 runs per day for 5 operating days on one LIAISON^®^ XL Analyzer (N=90).

A cross-reactivity study was performed to evaluate: 1) other SARS viruses (HCoV-229E, HCoV-HKU1, HCoV-OC43, and HCoV-untyped); 2) conditions caused by other viruses that may cause symptoms similar to SARS-CoV-2 infection; 3) infectious diseases caused by other organisms; and 4) conditions that may result in atypical immune system activity. Samples for the evaluation were collected before October 2019, prior to the Covid-19 pandemic. In addition, samples with potentially interfering factors such as triglycerides, hemoglobin, bilirubin, cholesterol, acetaminophen, ibuprofen and biotin were assessed with the LIAISON^®^ SARS-CoV-2 S1/S2 IgG assay.

### SARS-CoV-2 microneutralization assay (NT assay)

A neutralizing assay described elsewhere was used to determine the neutralization titer against SARS-CoV-2 (Percivalle et al, submitted for publication). Briefly, 50 μl of diluted serum (4-fold serial dilutions from 1:10 to 1:640) were added to an equal volume of viral suspension (tissue culture infectious dose 50 of a SARS-CoV2 strain isolated from a symptomatic patient), incubated, and then combined with Vero-E6 cells. After incubation, the cells were stained with Gram’s crystal violet solution. Wells were scored to evaluate the degree of cytopathic effect compared to the virus control. Neutralizing titer was the maximum dilution evidencing a reduction of 90% of cytopathic effect. In this study a titer of ≥1:40 was considered positive. This test was used to confirm positive serological samples used in the clinical studies, and to determine the neutralization effectiveness of the samples for the identification of convalescent donors living in the first Italian Red Zone (Percivalle et al, submitted for publication).

### Clinical Samples

This observational study used de-identified fresh or frozen residual samples collected between 2011 and April 2020 at the Policlinico San Matteo in Pavia, and at the Niguarda Hospital in Milan (Italy). Sample groups included: 1) paired samples (admission and discharge) from patients affected by Covid-19 and hospitalized with moderate symptoms (confirmed by RT-PCR, N=31); 2) sets of samples (admission and follow-ups) from patients affected by Covid-19 and hospitalized in the ICU with severe symptoms (confirmed by RT-PCR, N=16); 3) samples from patients affected by Covid-19 and hospitalized in ICU with severe symptoms (confirmed by RT-PCR, N=21); 4) samples from patients affected by Covid-19 testing positive by RT-PCR (N=37); 5) samples from subjects collected before the outbreak of Covid-19 (lab routine 2011, N=1140); 6) samples from subjects not infected by SARS-CoV-2 but affected by other coronaviruses, i.e. HCoV-229E, HCoV-HKU1, HCoV-OC43, or HCoV-untyped strain (N=10); 7) samples from subjects testing negative by RT-PCR (N=50); 8) samples negative by the SARS-CoV-2 NT assay described in Materials & Methods, collected from subjects during the outbreak (N=180); and 9) NT assay-positive samples collected from subjects during the outbreak (N=124).

Pertinent additional information included sample collection time, days from diagnosis (hospital or ICU admission), and severity of symptoms (mild, moderate, severe). The protocol for this study (de-identified remainders) was determined to be exempt under existing ethics committee regulations.

### Diagnosis

Diagnosis was based on results from routine RT-PCR used in the clinical evaluation (25), as well as NT-assay results. Samples not infected by SARS-CoV-2, but affected by other coronaviruses, were classified by sequencing.

### Statistical analyses

The statistical program R 3.5 and MedCalc 19.2 were utilized for all analyses presented. A preliminary exploration using Box-Cox methodology suggested that statistical analyses be done on the logarithmic scale due to skewed distribution of the measurements. Data supporting Figures and Tables in this manuscript will be made fully available upon request.

## ACKNOWLEDGMENTS

We acknowledge the full financial support from DiaSorin S.p.A. for this clinical trial. Employees of DiaSorin participated in the study design, data collection and interpretation, and in the preparation of the manuscript.

We thank the product development team at DiaSorin (Andrea Dal Corso, Lorenzo Querin, Elisa Mazzoleni, Piernatale Brusasca, Elisa Ghezzi, Clara Rossini, Valeria Rigamonti, Massimo Panizzo, Paolo Ingallinella, Davide Zanin), all the clinical coordinators for sample collection and analysis, data collection and interpretation, as well as Prof Douglas Hawkins for assistance with the statistical analyses.

Conflict of interest statement: FB, CZ, MC, FAB, LP and AC are employees of DiaSorin the manufacturer of the LIAISON^®^ SARS-CoV-2 S1/S2 IgG test. CFP spoke for, participated at boards of, and/or received research grants from Abbvie, Janssen, Cepheid, Viiv, Gilead, Merck, Lilly, DiaSorin, Elitech. FB spoke at sponsored meetings, participated at boards of, and/or received research grants from Abbott, Qiagen, Roche, Takeda, Viiv, Merck, DiaSorin, Elitech, HuMabs, Biotest, ABL, NTP. AS, AB, CV,DC, EP, and APD have no conflicts of interest to disclose.

## ABBREVIATIONS

ICU: intensive care unit
NT assay: neutralization assay
PPA: positive predictive agreement
PPV: positive predictive value
NPV: negative predictive value
S1: Spike protein fragment 1
S2: Spike protein fragment 2
IgG: immunoglobulin G

